# Extraintestinal roles of intestinal vitamin D receptor in protecting against dysbiosis and tumorigenesis in breast cancer

**DOI:** 10.1101/2022.05.17.492300

**Authors:** Yong-Guo Zhang, Jilei Zhang, Shreya Deb, Shari Garrett, Yinglin Xia, Jun Sun

**Affiliations:** Division of Gastroenterology and Hepatology, Department of Medicine, University of Illinois at Chicago, IL 60612, USA; Department of Microbiology and Immunology, Department of Medicine, University of Illinois at Chicago, IL 60612, USA; UIC Cancer Center, University of Illinois at Chicago, IL 60612, USA; Jesse Brown VA Medical Center Chicago, IL (537), USA

**Keywords:** Dysbiosis, breast cancer, barrier function, butyrate-producing bacteria, butyrate, inflammation, gut-breast-axis, *Lactobacillus plantarum*, probiotics, tight junctions, VDR

## Abstract

The microbiota play critical roles in regulating the function and health of intestine and extraintestinal organs. A fundamental question is whether there is an intestinal-microbiome-breast axis during the development of breast cancer. If yes, what are the roles of host factors? Vitamin D receptor (VDR) involves host factors and the human microbiome. *Vdr* gene variation shapes the human microbiome and VDR deficiency leads to dysbiosis. We hypothesized that intestinal VDR protects hosts against tumorigenesis in breast. Reduced VDR mRNA expression was observed in patients with breast cancer. We used a 7,12-dimethylbenzanthracene (DMBA)-induced breast cancer model in intestinal epithelial VDR knockout (VDR^ΔIEC^) mice. We reported that VDR^ΔIEC^ mice with dysbiosis are more susceptible to breast cancer induced by DMBA. Intestinal and breast microbiota analysis showed that lacking VDR leads to bacterial profile shift from normal to susceptible carcinogenesis. We found enhanced bacterial staining within breast tumors. At the molecular and cellular levels, we identified the mechanisms by which intestinal epithelial VDR deficiency led to increased gut permeability, disrupted tight junctions, microbial translocation, and enhanced inflammation, thus increasing the tumor size and number in breast. Furthermore, treatment with beneficial bacterial metabolite butyrate or probiotic *Lactobacillus plantarum* reduced the breast tumors, enhanced the tight junctions, and inhibited inflammation in the VDR^ΔIEC^ mice. Gut microbiome contribute to the pathogenesis of diseases, not only in the intestine, but also in the breast. Our study provides new insights into the mechanism by which intestinal VDR dysfunction and gut dysbiosis led to high risk of extraintestinal tumorigenesis. Gut-tumor-microbiome interactions indicate a new target in the prevention and treatment of breast cancer.

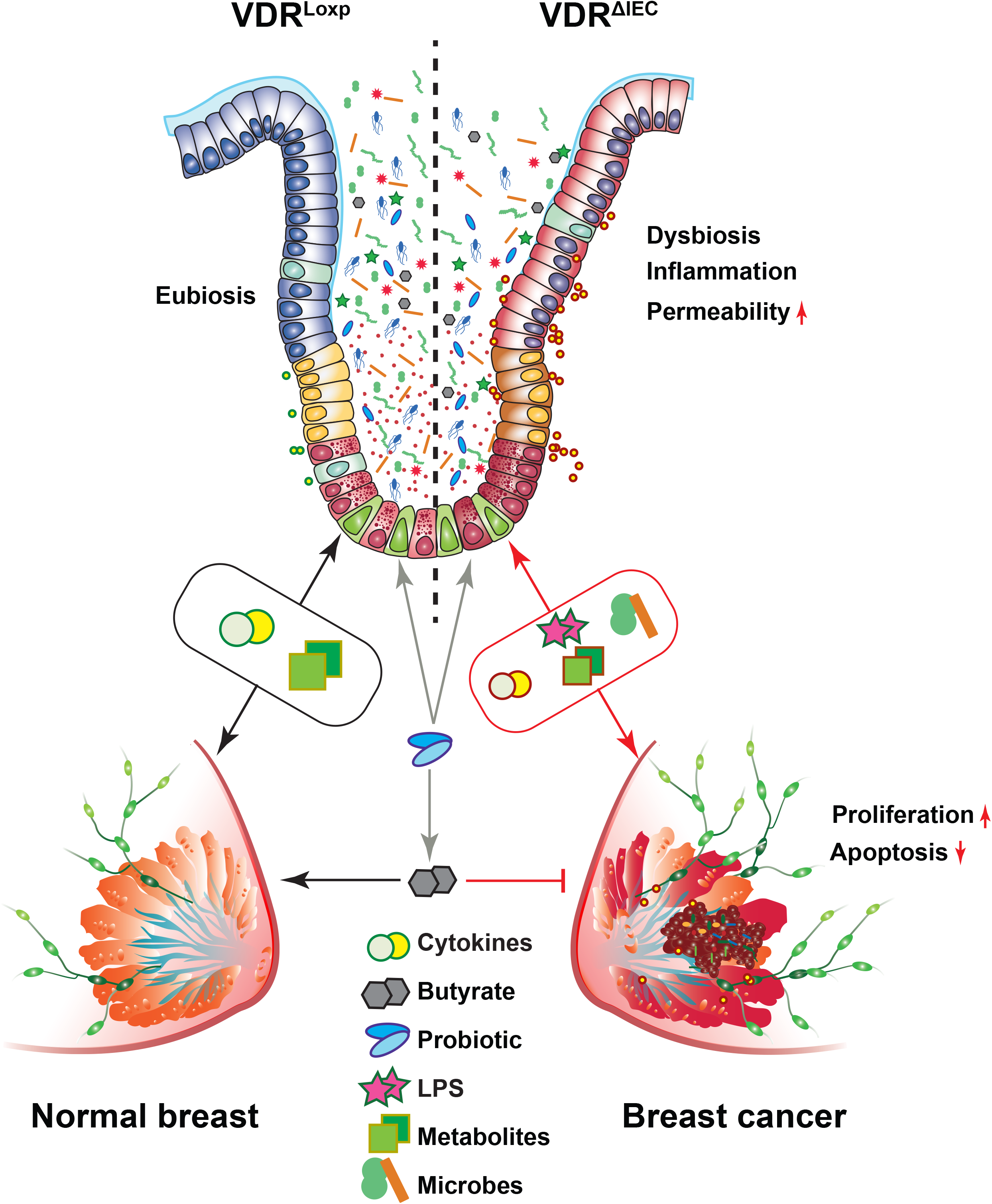

## Introduction

Vitamin D is a group of fat-soluble steroids responsible for multiple biological effects. The active form of vitamin D, in conjunction with its own receptor VDR, exerts important roles in modulating both mucosal immunity and normal growth of epithelia cells ^1, 2^. The dysregulation of the vitamin D/VDR is known to increase the risk of various human disorders ^3, 4, 5^. The parallel appreciation of a role for VDR in cancer biology began approximately 3 decades ago and has subsequently increased in the understanding of its actions in normal and malignant systems ^6^.

The VDR dependent regulation of the gut microbiome in human and animal studies represents a newly identified and highly significant role for VDR ^7, 8^. We have demonstrated that the variations of human *Vdr* gene shape the gut microbiome and VDR deletion leads to dysbiosis ^8^. Our studies support the critical role of VDR in maintaining intestinal and microbial homeostasis ^9-11^. We established the first conditional deletion of intestinal epithelial VDR mouse model (VDR^ΔIEC^) and demonstrated that intestinal bacterial abundance and function are significantly altered in VDR^ΔIEC^ mice ^7, 12^. VDR^ΔIEC^ mice were also susceptible to inflammatory triggers ^7^, indicating that intestinal VDR contributes to host protection against injury and inflammation. Dysbiosis and chronic inflammation are important contributors to the cancer development.

Research progress on the VDR-microbiome has established a microorganism-induced program of epithelial cell homeostasis and repair in the intestine ^13^. Dysregulation of bacterial-host interaction can result in chronic inflammatory and over-exuberant repair responses in the development of cancer ^14-19^. Vitamin D deficiency due to lack of sun exposure, especially in the region with special living habits, is considered an important cause of breast cancer ^20^. Polymorphisms of *vdr* gene (Bsm1, Apa1, Fok1, and Poly(A)) were reported to increase susceptibility of breast cancer ^21, 22^. Even though vitamin D/VDR is an active topic in cancer research, the mechanism underlying host-microbiome interactions in tumorigenesis is incompletely understood ^10, 11, 23, 24^. We know little about the effects and mechanisms by which intestinal epithelial VDR and microbiome influence dysbiosis in breast cancer.

In the current study, we hypothesized that intestinal VDR protects hosts against tumorigenesis in breast. Reduced VDR mRNA expression was observed in breast samples from patients with breast cancer. We found that VDR^ΔIEC^ mice with intestinal dysbiosis are more susceptible to breast cancer induced by DMBA. Intestinal and breast microbiota analysis showed that lacking VDR leads to bacterial profile shift from normal to susceptible carcinogenesis. At the cellular levels, we identified the mechanisms by which intestinal epithelial VDR deficiency led to increased gut permeability, disrupted tight junctions, microbial translocation, and enhanced inflammation, thus increasing the tumor size and number in breast. Furthermore, treatment with beneficial bacterial metabolite butyrate or probiotics reduced the breast tumors in the VDR^ΔIEC^ mice. Gut dysbiosis contribute to the pathogenesis of diseases, not only in the intestine, but also in the breast. Our study provides new insights into the mechanism by which intestinal VDR dysfunction and gut dysbiosis led to high risk of extraintestinal tumorigenesis in the breast.

## Results

### Downregulated VDR expression in breast cancer patients and altered bacterial diversity and functions in VDR deficient mice

We first examined the expression of the *vdr* gene in normal and breast cancer patient samples by reviewing the GEO database GSE 7904 ^25^. Reduced VDR mRNA expression was observed in breast samples from breast cancer patients (**Figure. 1a**). We then checked whether intestinal VDR deletion has any effects on the microbiome and the risk of breast cancer. We performed the metagenomic sequencing of 20 fecal samples from 2 groups, the group of VDR^ΔIEC^ mice, which conditionally deleted VDR from the intestinal epithelial cells (5 males and 5 females), and the control group of VDR^loxp^ mice (3 males and 7 females). After removing the repeated sequence with ≥99% identity, 52,892,651 taxonomic alignments with an average of 2,644,633 reads per sample, and 27,893,990 functional alignments with an average of 1,394,700 reads per sample were generated.

**Figure 1.**
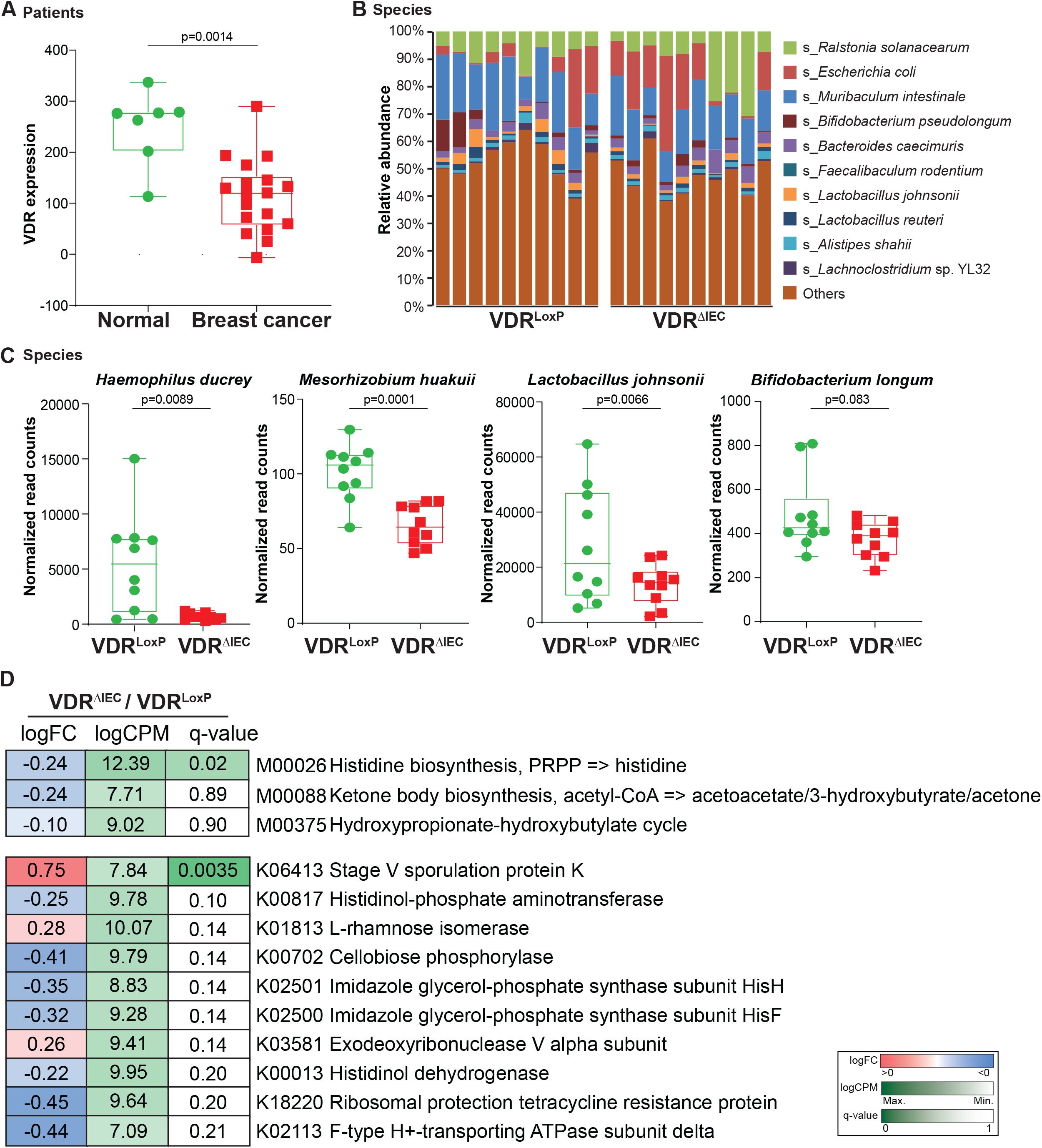
Reduced VDR expression in breast cancer patients and altered taxonomic community of intestinal bacteria in VDR^ΔIEC^ mice compared with VDR^loxp^ mice. **(a)** Reduced VDR expression in breast biopsy samples from breast cancer patients (GEO database 7904). Data are expressed as mean ± SD; Normal, nL=L7; Breast cancer, nL=L18; Welch’s *t*-test. **(b)** Relative bacterial abundance at the species level is shown with the top 10 species, and less abundant species were grouped as “others”. Each bar represents an individual mouse, n=10 per group. **(c)** The presentive bacteria species that were marked altered after VDR conditionally deletion. Data are expressed as mean ± SD, Welch’s *t*-test, n=10 each group. **(d)** Differential analysis of functional genes in feces of conditional VDR-knockout mice. The KEGG MODULE database consists of KEGG modules identified by M numbers, which are manually defined functional units of gene sets. The KEGG Module ortholog table is a useful tool to check completeness and consistency of genome annotations. It shows currently annotated genes in individual genomes for a given set of K number. The items with q-value ≤0.05 in pairwise comparisons or butyrate-related were selected. The fold-change (log_2_FC), counts per million (log_2_CPM), and q-value were colored using the key as indicated on the right side of the figure, n=10 each group. All *p* values are shown in the figures.

As shown in **Fig.1b**, the bacterial community profiles had differences in diversity and compositions of the studied animals between the control group VDR^loxp^ mice and VDR^ΔIEC^. The top 10 most prevalent bacterial species in each animal were present individually, **i.e**. *Ralstonia solanacearum, Escherichia coli, Muribaculum intestinale, Bifidobacterium pseudolongum, Bacteroides caecimuris, Faecalibaculum rodentium, Lactobacillus johnsonii, Lactobacillus reuteri, Alistipes shahii*, and *Lachnoclostridium* sp. YL32 **(Figure 1b)**. *Haemophilus ducreyi* is a gram-negative bacterium and causative agent of genital ulcer disease chancroid ^26^, and was found significantly downregulated in the VDR^ΔIEC^ mice **(Figure 1c)**. *Mesorhizobiu*m *huakuii* induces the formation of nitrogen-fixation nodules on its host plant *Astragalus sinicus* and has been assigned to a new biovariant ^27^. *M. huakuii* isolates were also found to have endotoxic activity against lipopolysaccharides ^28^ and were significantly downregulated in our VDR^ΔIEC^ mice. Moreover, there are two beneficiary bacteria species, *Lactobacillus johnsonii* and *Bifidobacterium longum*, were found markedly downregulated in the VDR^ΔIEC^ mice **(Figure 1c)**.

In addition to bacterial diversity and abundance, shotgun metagenomic sequencing could reveal microbial functional alterations through functional analysis. We performed taxonomic functional module, pathway, and table analyses, which were mainly based on the set of related biosynthesis, individual pathway, and functional genes, respectively. Then, we performed differential analysis on the functional profiling to show the impacts of VDR conditional deletion on these identified functions in modules, pathways, and tables. The histidine biosynthesis pathway, which is an ancient metabolic pathway present in bacteria, archaea, lower eukaryotes and plants and has a fundamental regulatory processes in bacteria (e.g., proton buffering and metal ion chelation) ^29, 30^, was found significantly (q<0.05) downregulated in the VDR^ΔIEC^ mice but upregulated in other pairwise comparisons without any statistical difference **(Figure 1d)**. Interestingly, some butyrate-related modules were also downregulated in the VDR^ΔIEC,^ mice, and the gene encoding stage V sporulation protein K, which is essential for sporulation and specific to stage V sporulation ^31^, was significantly (q<0.01) upregulated in the VDR^ΔIEC^ mice **(Figure 1d)**.

### VDR^ΔIEC^ mice developed more and larger breast tumors, compared to the VDR^loxp^ mice

Gut dysbiosis is associated with the development of breast cancer ^32-36^. Our metagenomic sequencing results indicated intestinal epithelial VDR knock out induced gut microbiome dysbiosis. We then investigated the role of intestinal VDR in the development of breast cancer, using a DMBA mouse model (**Figure 2a**). Chemically induced rodent models of breast cancer have been extensively used to reflect the initiation and progression of human breast cancer ^37-40^. We found a striking difference in breast tumor incidence between the VDR^loxp^ and VDR^ΔIEC^ mice with DMBA treatment. Representative breast tumors were shown in **Figure 2b**. The number of breast tumors was significantly increased in the VDR^ΔIEC^ mice, compared with the VDR^loxp^ mice (**Figure 2c**). The volumes of the tumors were significantly larger in the VDR^ΔIEC^ mice, compared with the VDR^loxp^ mice (**Figure 2d**). Furthermore, pathological analysis of breast samples indicated the tumor size differences between VDR^loxp^ and VDR^ΔIEC^ mice DMBA experimental groups **(Figure 2e)**. However, intestinal morphology did not change in VDR^loxp^ and VDR^ΔIEC^ mice with DMBA treatment (**Figure 2f and 2g**).

**Figure 2.**
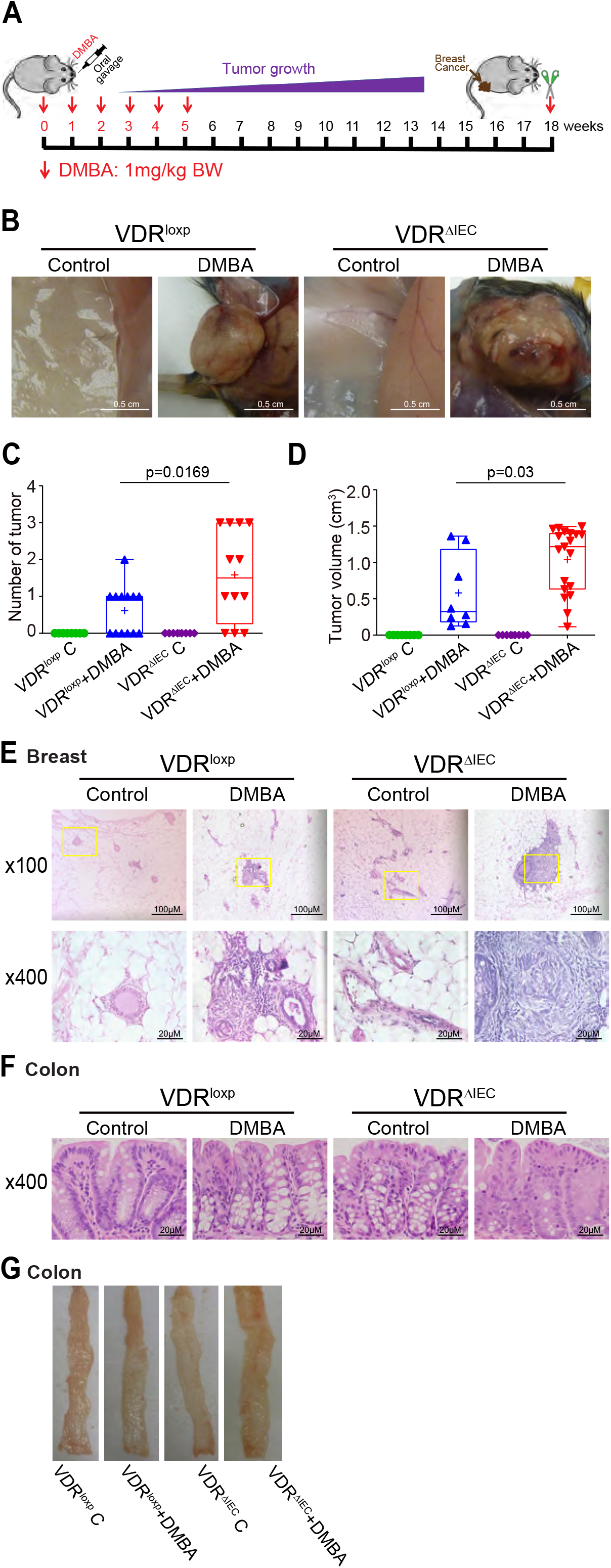
VDR^ΔIEC^ mice developed larger and more breast tumors. **(a)** The schematic overview of DMBA-induced breast cancer model. Mice were given 1.0 mg of DMBA in 0.2 ml of corn oil by oral gavage once a week for 6 weeks. The samples were harvested at week 18. **(b)** The breast tumors in situ. Representative mammary glands from different groups. **(c)** The number of breast tumors significantly increased in the VDR^ΔIEC^ mice compared with the VDR^loxp^ mice. Data are expressed as mean ± SD. N = 8-13, one-way ANOVA test. **(d)** The breast tumor volumes were significantly bigger in size in the VDR^ΔIEC^ mice compared with the VDR^loxp^ mice. Data are expressed as mean ± SD. n = 8-13, one-way ANOVA test. **(e)** Representative H&E staining of mammary glands from the indicated groups. Images were from a single experiment and are representative of 8-13 mice per group. **(f)** Representative H&E staining of intestinal from the indicated groups. Images were from a single experiment and are representative of 8-13 mice per group. All *p* values were shown in the figures. **(g)** Representative photographs of colons from the indicated groups.

### Intestinal epithelial VDR deletion led to decreased VDR expression, increased proliferation, and decreased apoptosis in breast tumor tissues

In the VDR^ΔIEC^ mice, we observed significant downregulation of VDR at the protein level in the breast tumor tissue **(Figure. 3a)**. Reduced VDR expression was confirmed via IHC staining of breast tissues of the control and DMBA treated VDR^loxp^ and VDR^ΔIEC^ mice **(Figure. 3b)**. Our WB and IHC data of proliferative marker p-β-catenin (ser 552) ^41-43^ showed that p-β-catenin (ser552) in the breast tumor tissue was significantly increased in the VDR^ΔIEC^ mice, compared to the VDR^loxp^ mice **(Figure. 3a and 3c)**. Apoptosis positive cells were decreased in breast tumor tissue of VDR^ΔIEC^ mice, compared with VDR^loxp^ mice by TUNEL staining **(Figure 3d)**. Altered cell proliferation and apoptosis in the breast of VDR^ΔIEC^ mice ultimately enhanced its susceptibility to carcinogenesis.

**Figure 3.**
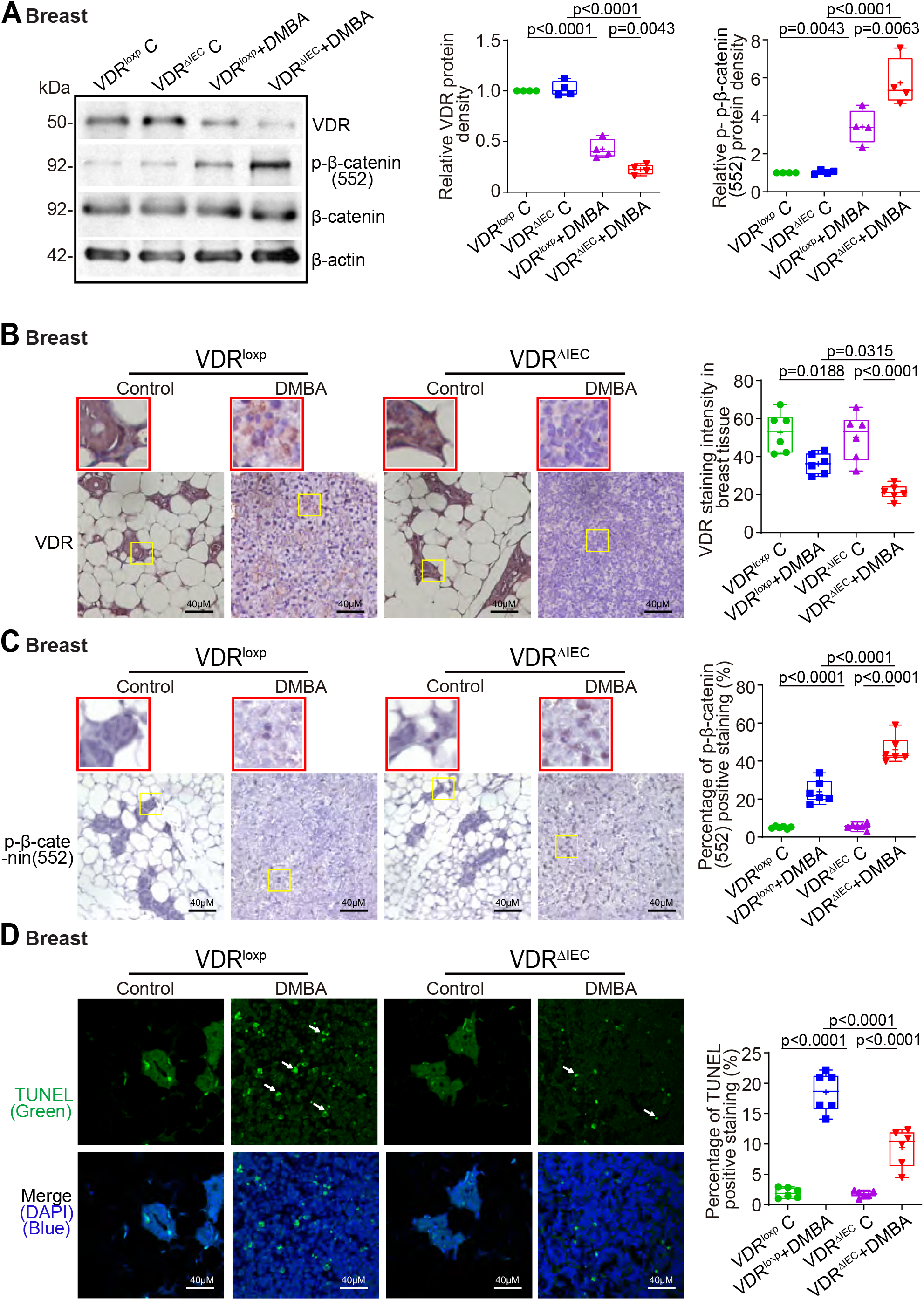
Intestinal epithelial VDR deletion led to decreased VDR expression, increased proliferation, and decreased apoptosis in breast tumor tissues. **(a)** Decreased VDR protein expression and increased p-β-catenin (Ser552) expression in mammary glands tumors of VDR^ΔIEC^ mice, compared with VDR^loxp^ mice. Data are expressed as mean ± SD. N = 4, one-way ANOVA test. **(b)** VDR was decreased in breast tumors of VDR^ΔIEC^ mice compared with VDR^loxp^ mice by IHC staining. Images are from a single experiment and are representative of 6 mice per group. Red boxes indicate the selected area in higher magnification. **(c)** p-β-catenin (ser552) expression increased in breast tissues of VDR^ΔIEC^ mice compared with VDR^loxp^ mice by IHC staining. Images are from a single experiment and are representative of 6 mice per group. Red boxes indicate the selected area in higher magnification. **(d)** Apoptosis positive cells were decreased in breast tissue of VDR^ΔIEC^ mice compared with VDR^loxp^ mice by TUNEL staining. Images are from a single experiment and are representative of 6 mice per group. All *p* values are shown in the figures.

### Lacking intestinal VDR led to increased gut permeability, disrupted tight junctions, microbial translocation, and enhanced inflammation

Gut dysbiosis usually increases harmful intestinal bacteria, which may release more enterotoxins e.g. LPS, damaging tight junctions (TJs) in epithelial cells, thereby increasing the permeability of the intestine and elevate the risk for cancer ^32, 44^. To test intestinal permeability, mice were gavaged with Fluorescein Dextran. Four hours later, blood samples were collected for fluorescence intensity measurement. Higher fluorescence intensity indicated increased intestinal permeability. As shown in **Figure. 4a**, DMBA treatment increased intestinal permeability in both VDR^loxp^ and VDR^ΔIEC^ mice, while the VDR^ΔIEC^ mice exhibited significantly higher permeability post treatment. Based on the *in vivo* intestinal permeability data, we hypothesized that the TJ proteins might be altered in the DMBA treated mice. In the VDR^ΔIEC^ mice, we observed significant downregulation of ZO-1 at the protein level in the colon **(Figure. 4b)**. Reduced and disorganized ZO-1 was confirmed by the immunostaining of colon in the DMBA treated mice **(Figure. 4c)**. Butyrate synthesis by anaerobic bacteria can occur via butyryl-coenzyme A (CoA): acetate CoA-transferase ^45^. We found that intestinal VDR deficiency led to dysbiosis and shift of bacterial profile. Expression of butyryl-CoA transferase decreased in the feces of the VDR^ΔIEC^ mice, compared to VDR^loxp^ mice. *E. coli* was enhanced in VDR^ΔIEC^ mice, compared to those in VDR^loxp^ mice **(Figure. 4d)**. Chronic inflammation is one of the key factors that contribute to breast cancer. We found that serum LPS, cytokines IL-1β, IL-6, IL-5 and TNF-α were significantly higher in the DMBA treated VDR^ΔIEC^ mice, compared to the VDR^loxp^ mice **(Figure. 4e)**. More universal bacteria were found in breast tumors of VDR^ΔIEC^ mice by FISH staining **(Figure. 4**f), suggesting the enhanced local bacteria within the breast.

**Figure 4.**
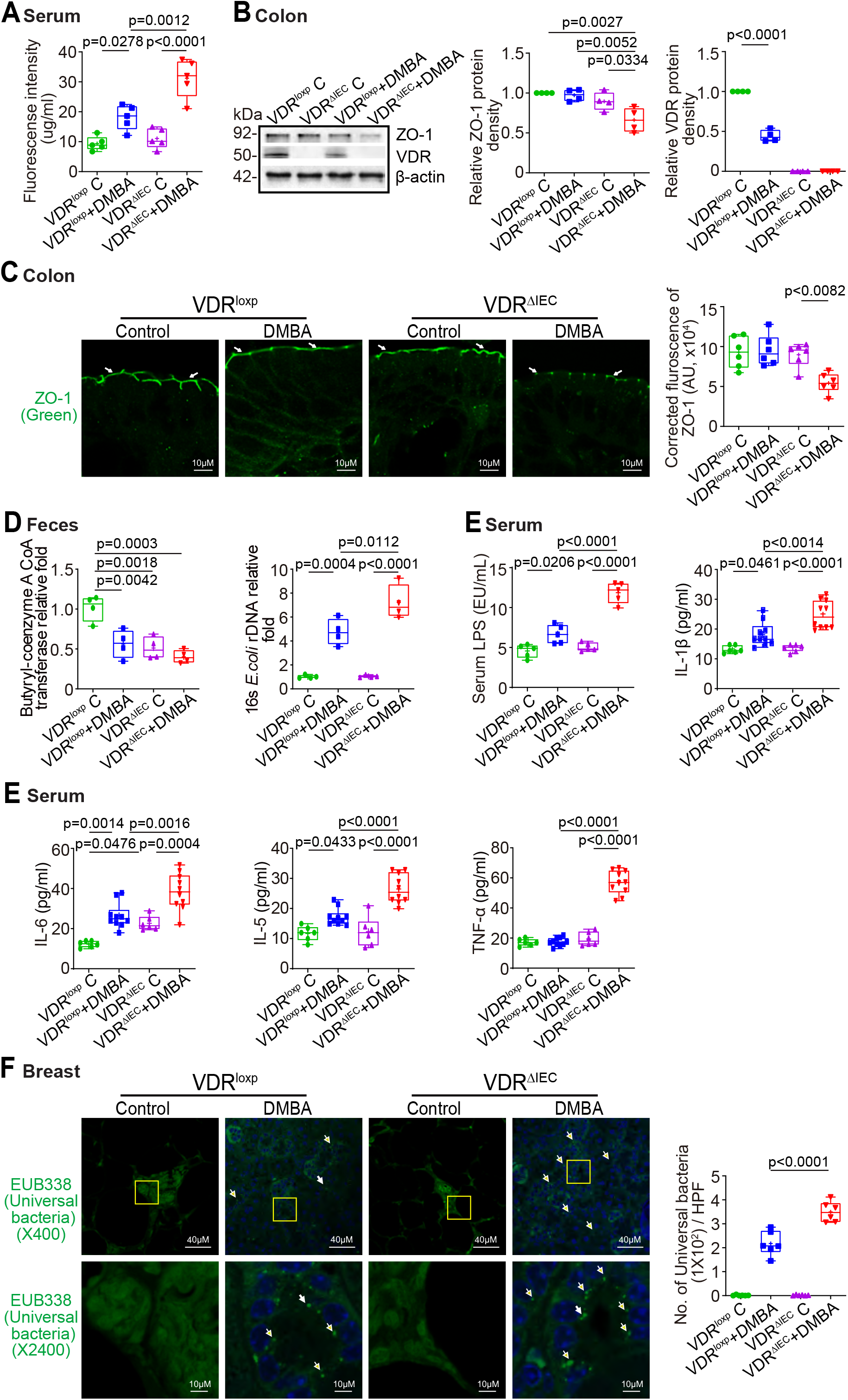
Increased intestinal permeability, decreased ZO-1 expression, chronic inflammation and increased universal bacteria in breast tumors of VDR^ΔIEC^ mice compared with VDR^loxp^ mice. **(a)** Intestinal permeability increased in the DMBA-induced VDR^ΔIEC^ breast cancer model. Fluorescein Dextran (Molecular weight 3 kDa, diluted in HBSS) was gavaged (50 mg/g mouse). Four hours later, mouse blood samples were collected for fluorescence intensity measurement. Data are expressed as mean ± SD. N = 5, one-way ANOVA test. **(b)** ZO-1 expression decreased in the intestine of VDR^ΔIEC^ mice after DMBA treatment, compared with VDR^loxp^ mice. Data are expressed as mean ± SD; n = 4, one-way ANOVA test. **(c)** ZO-1 expression decreased in intestinal VDR^ΔIEC^ mice after DMBA treatment compared with VDR^loxp^ mice by IF staining. Images are from a single experiment and are representative of 6 mice per group. Data are expressed as mean ± SD. N = 6, one-way ANOVA test. **(d)** Lacking intestinal VDR led to dysbiosis and shift of bacterial profile. Expression of Butyryl-coenzyme A CoA transferase decreased in control and tumors in VDR^ΔIEC^ mice compared to VDR^loxp^ mice. *E*.*coli* was enhanced in tumors in VDR^ΔIEC^ mice compared to VDR^loxp^ mice. Data are expressed as mean ± SD. N = 4, one-way ANOVA test. **(e)** Serum LPS, IL-1β, IL-6, IL-5, IL-6 and TNF-α were significantly higher in tumors in VDR^ΔIEC^ mice compared to VDR^loxp^ mice. Serum samples were collected from VDR^loxp^ and VDR^ΔIEC^ mice with or without tumor, then cytokines were detected by Luminex detection system. Data are expressed as mean ± SD. N = 6-10, one-way ANOVA test. **(f)** More universal bacteria in breast tumor tissue of VDR^ΔIEC^ mice were found by fluorescence in situ hybridization. Images are from a single experiment and are representative of 6 mice per group. Data are expressed as mean ± SD. N = 6, one-way ANOVA test. All *p* values are shown in the figures.

### Butyrate treatment reduced the breast tumor number, increased breast VDR expression, decreased proliferation, and increased apoptosis in VDR^ΔIEC^ mice

Because butyrate-producing bacteria and butyrate synthesis related transferase were decreased in tumors in VDR^ΔIEC^ mice compared to the VDR^loxp^ mice, we then hypothesized that butyrate treatment in mice could reduce the formation of breast tumors. Female VDR^loxp^ and VDR^ΔIEC^ mice were treated with 2.5% butyrate in the drinking water starting at the age of 6-7 weeks, followed by scarification at the age of 18 weeks. The breast tumor number was significantly decreased in the VDR^ΔIEC^ mice treated with butyrate **(Figure 5a)**. The breast tumor volume was significantly smaller in the VDR^ΔIEC^ mice treated with butyrate, compared to those without butyrate treatment (**Figure 5b)**. Pathological analysis showed that the mammary glands were smaller in sizes in the VDR^ΔIEC^ mice treated with butyrate **(Figure 5c)**. Increased protein expressions of VDR and reduced p-β-catenin (552) were observed in breast tumors of the VDR^ΔIEC^ mice with butyrate treatment **(Figure 5d)**. Increased VDR expression was confirmed by IHC staining of breast tumor tissue in the VDR^ΔIEC^ mice treated with butyrate **(Figure 5e)**. In the VDR^ΔIEC^ mice treated with butyrate, we found significantly reduced p-β-catenin (ser552) in the breast tumors **(Figure 5f)**. Apoptosis positive cells were also significantly increased in breast tumors of VDR^ΔIEC^ mice with butyrate treatment by TUNEL staining **(Figure 5g)**.

**Figure 5.**
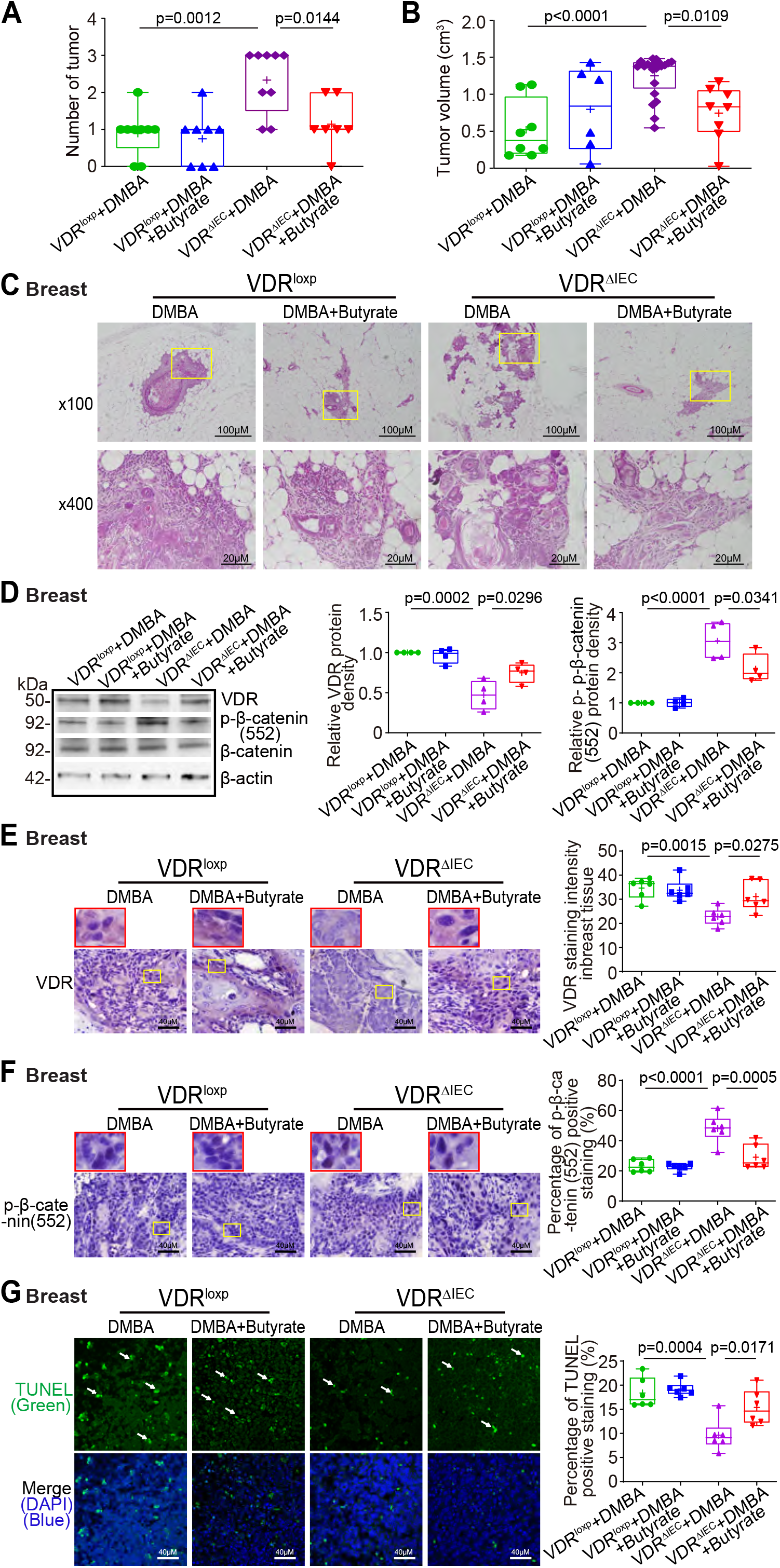
Butyrate treated VDR^ΔIEC^ mice had fewer and smaller tumors, increased breast VDR expression, decreased breast p-β-catenin (552) expression, and increased cell apoptosis. **(a)** The number of breast tumors significantly decreased in the VDR^ΔIEC^ mice with butyrate treatment mice. Data are expressed as mean ± SD. N = 7-9, one-way ANOVA test. **(b)** The breast tumor volumes were significantly smaller in the VDR^ΔIEC^ mice with butyrate treatment mice. Data are expressed as mean ± SD. N = 7-9, one-way ANOVA test. **(c)** Representative H&E staining of mammary glands from the indicated groups. Images are from a single experiment and are representative of 7-9 mice per group. **(d)** VDR expression increased, while p-β-catenin (Ser552) expression decreased in breast tumor tissue in the VDR^ΔIEC^ mice with butyrate treatment. Data are expressed as mean ± SD; N = 4, one-way ANOVA test. **(e)** VDR was increased in breast tumor tissue in the VDR^ΔIEC^ mice with butyrate treatment by IHC staining. Images are from a single experiment and are representative of 6 mice per group. Red boxes indicate the selected area in higher magnification. Data are expressed as mean ± SD. N = 6, one-way ANOVA test. **(f)** P-β-catenin (Ser552) expression decreased in breast tumor tissue in the VDR^ΔIEC^ mice with butyrate treatment by IHC staining. Images are from a single experiment and are representative of 6 mice per group. Red boxes indicate the selected area in higher magnification. Data are expressed as mean ± SD. N = 6, one-way ANOVA test. **(g)** Apoptosis positive cells were decreased in breast tumor tissue of VDR^ΔIEC^ mice with butyrate treatment by TUNEL staining. Images are from a single experiment and are representative of 6 mice per group. Data are expressed as mean ± SD. N = 6, one-way ANOVA test. All *p* values are shown in the figures.

### Butyrate treatment enhanced the intestinal TJs, corrected dysbiosis, and inhibited inflammation

We found that intestinal permeability was decreased in the VDR^ΔIEC^ mice with butyrate treatment (**Figure 6a)**. ZO-1 expression was increased in the colons of butyrate-treated VDR^ΔIEC^ mice **(Figure 6b)**. Increased ZO-1 expression was confirmed in the immunostaining of colon tissues of the VDR^ΔIEC^ mice with butyrate treatment **(Figure. 6c)**. There were increased butyryl-CoA transferase genes and decreased *E. coli* in the feces of VDR^ΔIEC^ mice with butyrate treatment **(Figure 6d)**. Serum enterotoxin LPS and proinflammatory cytokines (e.g. IL-1β, IL-5, IL-6, and TNF-α) were significantly lower in the VDR^ΔIEC^ mice with butyrate treatment **(Figure 6e)**, suggesting that butyrate protected mice from increased inflammation. Taken together, these data indicate that butyrate treatment led to a significantly reduced the breast tumors, enhanced the tight junctions, restored the microbiome, and inhibited inflammation in the VDR^ΔIEC^ mice.

**Figure 6.**
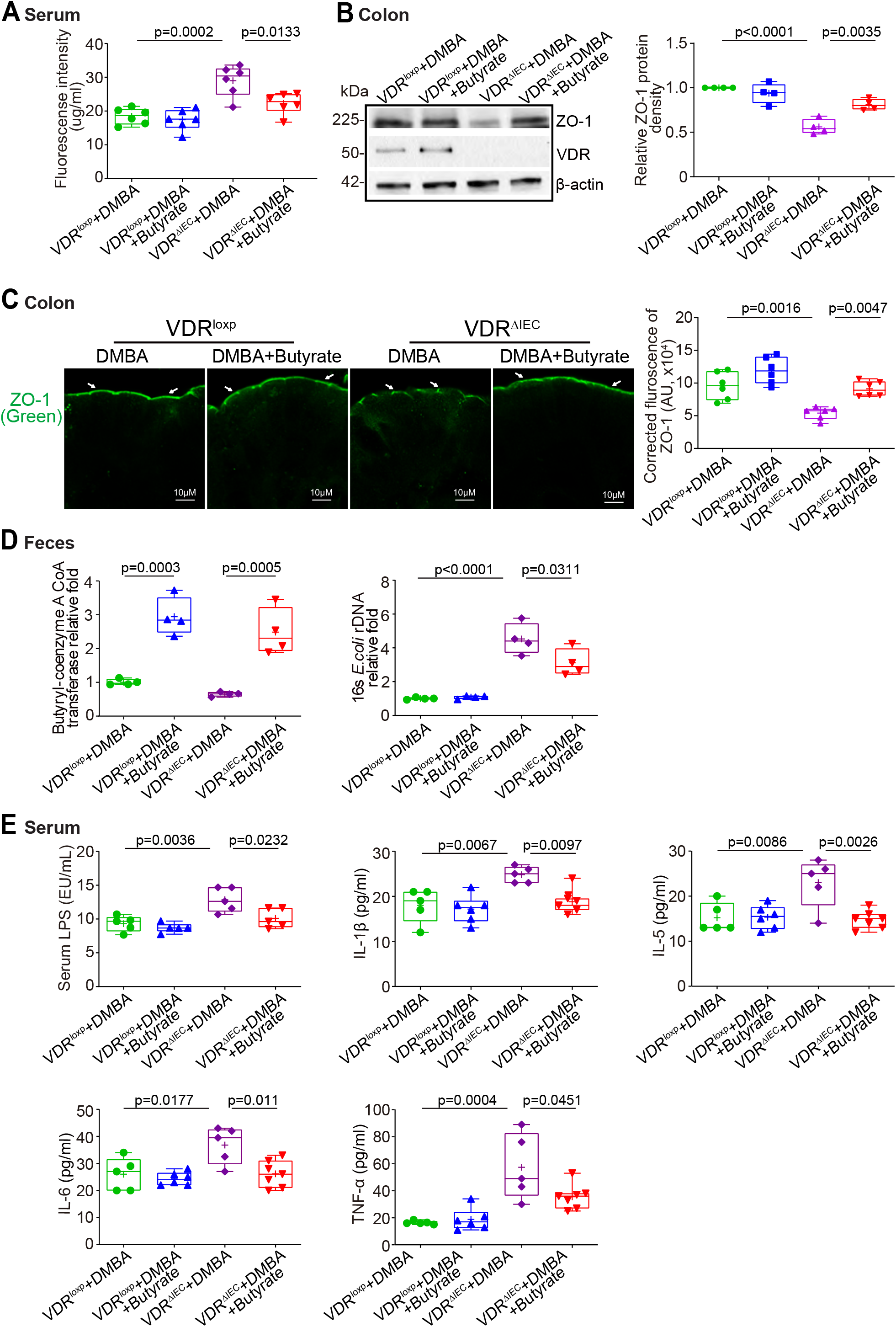
Butyrate treatment decreased intestinal permeability, increased intestinal ZO-1 expression, and decreased inflammation in the VDR^ΔIEC^ mice. **(a)** Intestinal permeability decreased in the VDR^ΔIEC^ mice with butyrate treatment. Data are expressed as mean ± SD. N = 6, one-way ANOVA test. **(b)** ZO-1 expression increased in the VDR^ΔIEC^ mice with butyrate treatment. Data are expressed as mean ± SD. N = 4, one-way ANOVA test. **(c)** ZO-1 expression increased in the VDR^ΔIEC^ mice with butyrate treatment by using IF staining. Images are from a single experiment and are representative of 6 mice per group. Data are expressed as mean ± SD. N = 6, one-way ANOVA test. **(d)** Butyrate treatment increased Butyryl-coenzyme A CoA transferase genes and decreased *E. coli* in the VDR^ΔIEC^ mice treated with butyrate. Data are expressed as mean ± SD. N = 4, one-way ANOVA test. **(e)** Butyrate treatment showed protection from increased inflammation in the VDR^ΔIEC^ mice. Serum LPS, IL-1β, IL-5, IL-6, and TNF-α were significantly lower in the VDR^ΔIEC^ mice treated with butyrate. Data are expressed as mean ± SD. N = 5-7, one-way ANOVA test. All *p* values are shown in the figures.

### Probiotics reduced the breast tumors, increased breast VDR expression, decreased proliferation, and increased apoptosis in VDR^ΔIEC^ mice

Probiotics *Lactobacillus plantarum* (*LP*) is known to increase VDR protein expression ^46^. To test the role of probiotics in breast cancer, female VDR^loxp^ and VDR^ΔIEC^ mice in DMBA-probiotics-treated groups were gavaged daily with *LP* starting at the 6-7 weeks of age, followed by scarification at the 18 weeks of age. The number of breast tumors significantly decreased in the VDR^ΔIEC^ mice with probiotic treatment mice **(Figure 7a)**. The breast tumors volumes were significantly smaller in the VDR^ΔIEC^ mice with probiotic treatment (**Figure 7b)**. Pathological analysis showed that mammary glands were smaller in size in the VDR^ΔIEC^ mice with probiotic treatment **(Figure 7c)**. In the probiotic-treated VDR^ΔIEC^ mice, we found that probiotic treatment significantly restored the expression of protein levels of VDR and reduced p-β-catenin (Ser552) in the breast tumors **(Figure 7d)**. Increased VDR expression was confirmed in the IHC staining of breast tumor tissues of VDR^ΔIEC^ mice with probiotic treatment **(Figure 7e)**. Decreased p-β-catenin^552^ expression was confirmed in IHC staining of breast tumor tissues of the VDR^ΔIEC^ mice with butyrate **(Figure 7f)**. By TUNEL staining, we found increased apoptotic cells in the VDR^ΔIEC^ mice with butyrate treatment **(Figure 7g)**.

**Figure 7.**
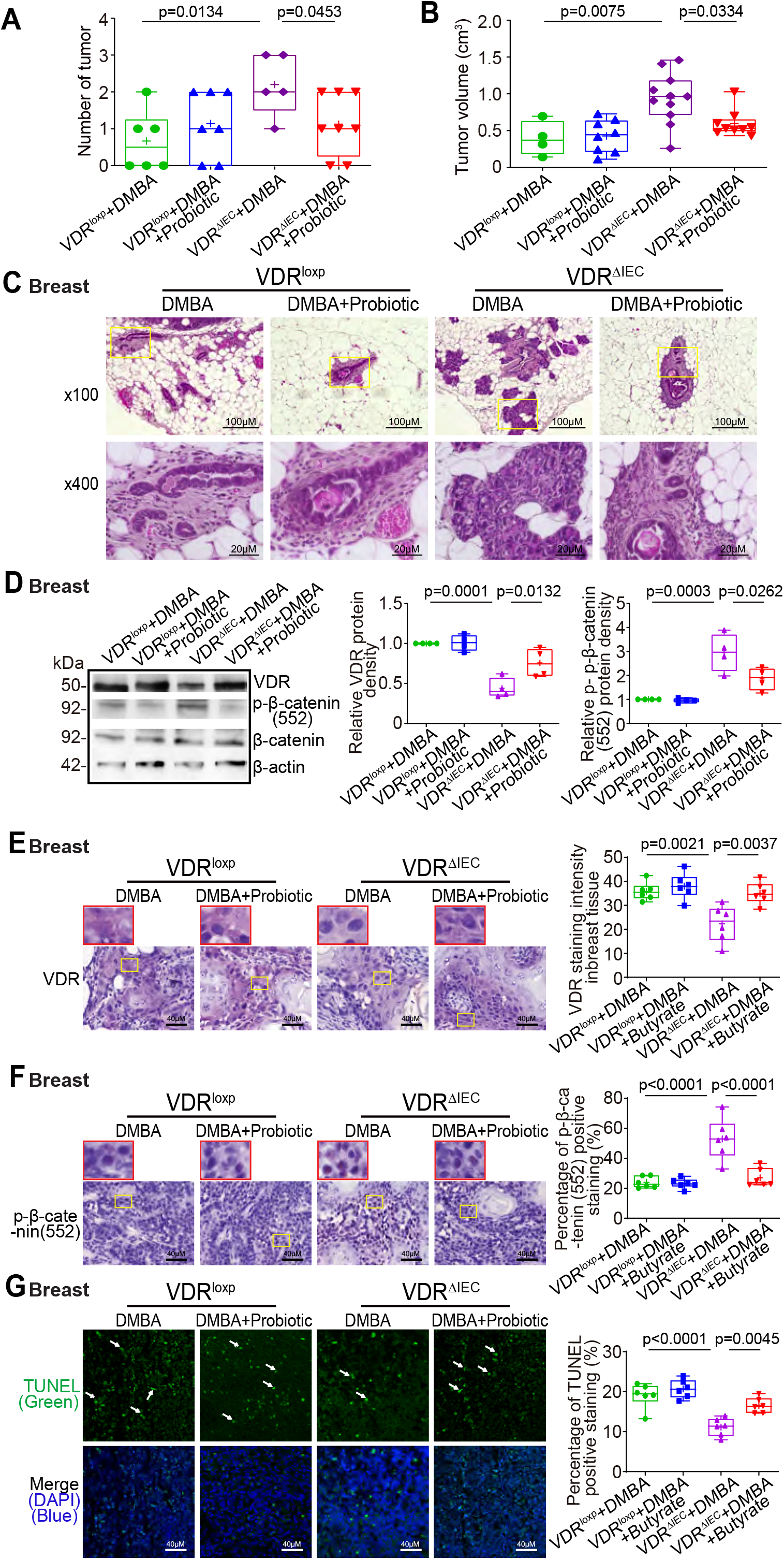
Probiotic-treated VDR^ΔIEC^ mice have fewer and smaller tumors, increased breast VDR expression, decreased expression of p-β-catenin (552), and increased cell apoptosis. **(a)** The number of breast tumors significantly decreased in the probiotics treated VDR^ΔIEC^ mice. Data are expressed as mean ± SD. N = 6-8, unpaired *t*-test. **(b)** The volume of breast tumors was significantly smaller in size in the probiotics treated VDR^ΔIEC^ mice. Data are expressed as mean ± SD. N = 6-8, one-way ANOVA test. **(c)** Representative H&E staining of mammary glands from the indicated groups. Images were from a single experiment and are representative of 6-8 mice per group. **(d)** VDR expression was increased, while p-β-catenin (552) expression was decreased in breast tumor tissue in the probiotic-treated VDR^ΔIEC^ mice. Data are expressed as mean ± SD. N = 4, one-way ANOVA test. **(e)** VDR was increased in breast tumor tissue in the VDR^ΔIEC^ mice with probiotic treatment by IHC staining. Images are from a single experiment and are representative of 6 mice per group. Red boxes indicate the selected area in higher magnification. Data are expressed as mean ± SD. N = 6, one-way ANOVA test. **(f)** P-β-catenin (552) expression decreased in breast tumor tissue in the VDR^ΔIEC^ mice with probiotic treatment by IHC staining. Images are from a single experiment and are representative of 6 mice per group. Red boxes indicate the selected area in higher magnification. Data are expressed as mean ± SD. N = 6, one-way ANOVA test. **(g)** Apoptosis positive cells were decreased in breast tumors of VDR^ΔIEC^ mice with probiotic treatment by TUNEL staining. Images are from a single experiment and are representative of 6 mice per group. Data are expressed as mean ± SD. N = 6, one-way ANOVA test. All *p* values are shown in the figures.

### Probiotics enhanced the intestinal TJs and reduced inflammation in the VDR^ΔIEC^ mice

We also found that the intestinal permeability decreased in the VDR^ΔIEC^ mice with probiotics treatment (**Figure 8a)**. ZO-1 expression increased in the colons of VDR^ΔIEC^ mice with probiotic treatment **(Figure 8b)**. Increased colonic ZO-1 expression was confirmed by immunostaining in the VDR^ΔIEC^ mice with probiotic treatment **(Figure 8c)**. Increased butyryl-CoA transferase genes and decreased *E. coli* were also observed in the VDR^ΔIEC^ mice with probiotic treatment **(Figure 8d)**. Moreover, serum LPS, IL-1β, IL-5, IL-6, and TNF-α were significantly lower in the VDR^ΔIEC^ mice treated with probiotics **(Figure 8e)**. These data suggested that probiotic treatment had several beneficial roles, e.g. reduced the breast tumors, enhanced TJs, and reduced inflammation, thus inhibiting tumorigenesis in the VDR^ΔIEC^ mice.

**Figure 8.**
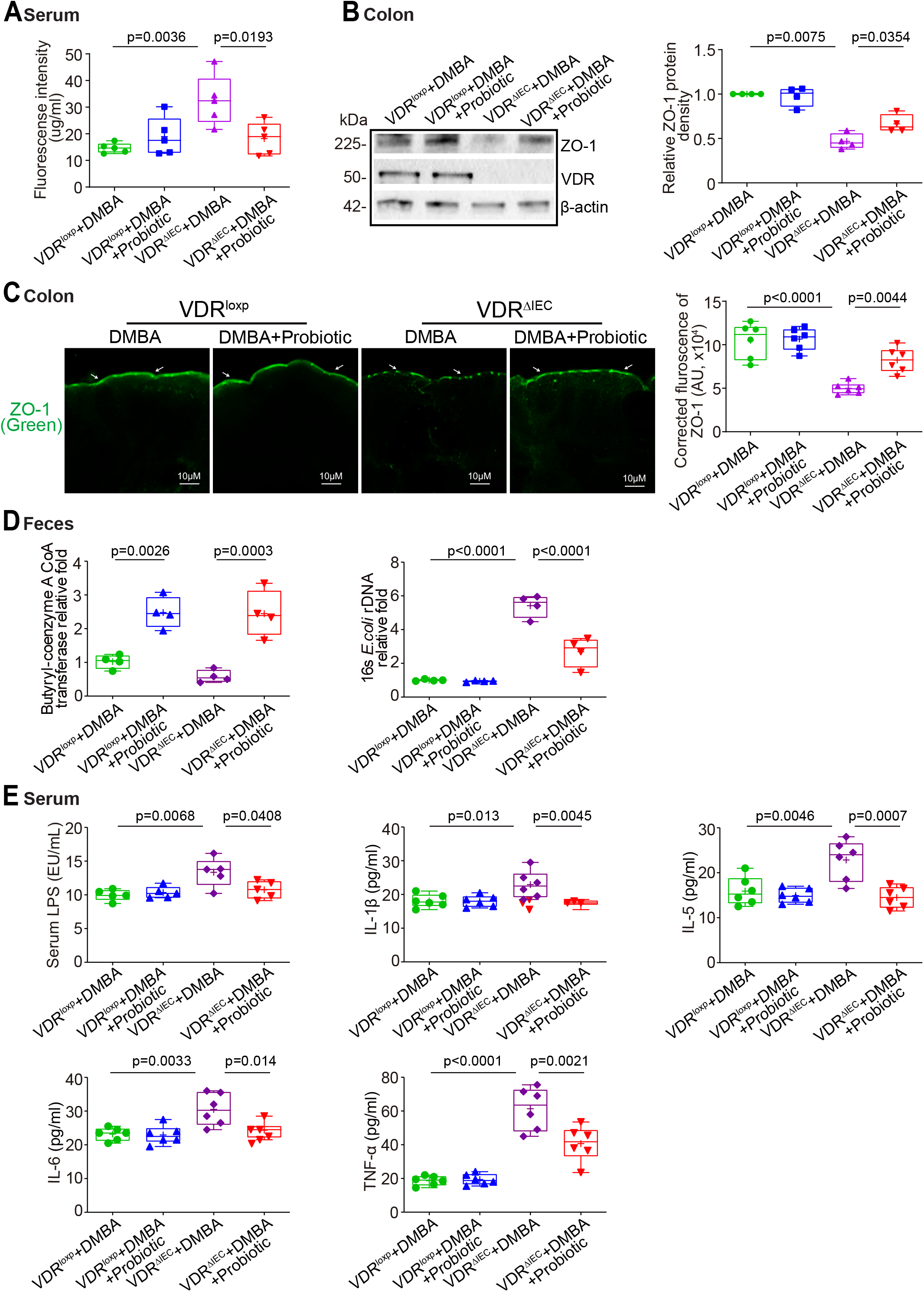
Probiotic-treated VDR^ΔIEC^ mice have decreased intestinal permeability, increased intestinal ZO-1 expression, corrected dysbiosis, and showed protection from increased inflammation. **(a)** Intestinal permeability decreased in the VDR^ΔIEC^ mice with probiotic treatment. Data are expressed as mean ± SD. N = 5, one-way ANOVA test. **(b)** ZO-1 expression increased in the VDR^ΔIEC^ mice with probiotic treatment. Data are expressed as mean ± SD. N = 4, one-way ANOVA test. **(c)** ZO-1 expression increased in the VDR^ΔIEC^ mice with probiotic treatment by the immunofluorescence staining. Images are from a single experiment and are representative of 6 mice per group. Data are expressed as mean ± SD. N = 6, one-way ANOVA test. **(d)** Probiotic treatment increased Butyryl-CoA transferase genes and decreased *E. coli* in the VDR^ΔIEC^ mice the. Data are expressed as mean ± SD. N = 4, one-way ANOVA test. **(e)** Probiotic treatment showed protection from increased inflammation in the VDR^ΔIEC^ mice. Serum LPS, IL-1β, IL-5, IL-6, and TNF-α were significantly lower in the VDR^ΔIEC^ mice treated with probiotics. Data are expressed as mean ± SD. N = 5-6, one-way ANOVA test. All *p* values are shown in the figures.

## Discussion

Our current study fills the gaps by revealing a previously unknown mechanism that intestinal VDR is important to normal host homeostasis and protects against breast cancer. Whereas the VDR^ΔIEC^ mice had normal VDR expression in the breast, the increased inflammation and bacterial loading/translocation still increased the risk of tumorigenesis in organs beyond the intestine. We identified increased intestinal permeability, chronic inflammation, and enhanced bacteria within the breast tumors. Furthermore, by manipulating the gut microbiome using a beneficial microbial metabolite or probiotic bacteria, we were able to reduce the tumor burden in the VDR^ΔIEC^ mice. Merging evidence has shown that enteric bacteria play a crucial role in the pathogenesis of breast cancer ^32, 47^. Our study further suggests new therapeutic targets for restoring intestinal VDR and microbiome functions in preventing breasr cancers (see Graphical Abstract). Clearly, research of intestinal VDR provides a framework to understand how the intestinal dysfunction may inadvertently promote the development of distant cancer. It is an extension of VDR’s normal role in defense and repair. These insights are important for understanding health as well as disease.

Multiple mechanisms describing VDR affects cancers have been found ^48^, but the specific relationship between the function of intestinal VDR and microbiome in breast tumorigenesis is not understood. An intricate symbiotic relationship has evolved between humans and microbes, especially the gut microbiota, which appear to influence the host at nearly every level and every organ system ^32, 47^. The microbiome, including the gut microbes and microbiota at the breast site ^49^, play an important role in breast health and diseases ^50^. However, little is known about the intestinal VDR regulation of the microbiome community in gut and breast. Our previous studies demonstrates that VDR KO mice are sensitive to bacterial invasion and exhibit severe damages in the intestine ^13 7 12^. VDR-bacterial interactions established an example of a microorganism-induced program of host homeostasis. Dysregulation of bacterial-host interactions can result in chronic inflammation and over-exuberant repair responses, and it is associated with the development of cancer ^14-19^, not just in the intestine, but also in other organs, e.g. the breast. Thus, our studies fill the gaps by investigating mechanisms by which intestinal epithelial VDR regulating breast microbiome and cancer.

Dysbiosis in the breast can change the microenvironment, which causes mastitis and poses a potential risk for breast cancer. A recent study has further shown that breast tumor-resident microbiota, albeit at low biomass, play an important role in promoting metastasis ^49^. Our study has shown the protective roles of lactic acid bacteria in breast tumorigenesis. Lactic acid bacteria are known for their beneficial health effects, including the anticarcinogenic feature. Different bacterial profiles in breast tissue exist between healthy women and those with breast cancer ^51^. Breast cancer patients had higher relative abundances of *Bacillus, Enterobacteriaceae* and *Staphylococcus*. Probiotic treatment helped to restore the healthy microbiome and their functions to produce beneficial metabolites. Our data support the idea to modulate the gut and mammary microbiome to reduce the risk of breast cancer.

Dysbiosis of the gut microbiome could indirectly participate in the development or progression of breast cancer via bacterial metabolites from the gut (SBA, SCFA, PAMPs, vitamin) and immune and inflammatory modulators (TLR, cytokines) that are inextricably interlinked with the microbiome from the GI tract ^32^. VDR expression increases epithelial integrity and attenuates inflammation ^52^. Interestingly, we found some butyrate-related modules were also downregulated in the VDR^ΔIEC^ mice. It has been reported that the butyrate can induce growth arrest, apoptosis or differentiation on breast cancer cell lines ^53^. We found that butyrate treatment in VDR^ΔIEC^ mice helped the hosts not only correct the dysbiosis, but also inhibit inflammatory cytokines, thus reducing breast tumors. Because low dose proinflammatory cytokines are sufficient to induce bacterial endocytosis by epithelial cells, sub-clinical or low-grade changes may tip the balance of tolerance towards full blown inflammation owing to subsequent intracellular microbial sensing and paracellular permeability damage. With the current knowledge of the microbiome in the development of various cancers, understanding of the interactions among epithelium, microbiome, and metabolites could help to develop strategies for managing chronic inflammation in diseases, including cancers.

Maintaining intestinal VDR functions and healthy gut microbiome will promote breast health. Our current study provides insights into alternative methods of enhancing VDR with bacterial product butyrate and probiotics, thus reducing the risk of breast tumors. Low VDR expression and diminished vitamin D/VDR signaling are observed in patients with breast cancer. Epidemiological and experimental studies have indicated a protective action of vitamin D against colorectal cancer ^54-60^. Vitamin D_3_ exerts its chemopreventive activity by interrupting a crosstalk between tumor epithelial cells and the tumor microenvironment in a VDR-dependent manner ^56^. Moreover, there is increasing interests in using vitamin D compounds for disease prevention and therapy^61^. Our findings advance our understanding of how VDR and the microbiome is involved into the pathogenesis of breast cancer. It also offers an additional avenue to treat breast cancer by restoring the function of intestinal VDR.

In conclusion, our study has demonstrated that intestinal epithelial VDR deficiency significantly influence intestinal barrier function, microbiome profile/location, and tumorigenesis in the breast. We provide definitive characterization of the intestinal epithelial VDR in shaping the microbiome in a breast cancer model. Gut-breast-microbiome interactions indicate a new target in preventing and treating breast cancer. It opens a new direction in the understanding of the microbial-VDR interactions in breast diseases and potential to develop a new protocol for risk assessment and prevention of extraintestinal illness.

## Materials and methods

### Human gene expression datasets

For expression analyses, we used microarray data reported in the NCBI Gene Expression Omnibus database ^62^. We gathered data by searching the Gene Expression Omnibus (https://www.ncbi.nlm.nih.gov/geo/) for expression profiling studies using breast biopsy samples from healthy controls as well as breast cancer biopsy samples from breast cancer subjects. We randomly selected the GEO database reference GSE 7904 ^25^, 7 healthy controls and 18 breast cancer patients were analyzed.

### Animals

The intestinal-specific VDR knockout VDR^ΔIEC^ mice were obtained by crossing the VDR^loxp^ mice with villin-cre mice (Jackson Laboratory, 004586), as we previously reported.^7, 63, 64^ We further backcrossed this strain with C57BL/6 mice for more than 10 generations after arriving our animal facility. Experiments were performed on 6- to 7-week-old female mice. Mice were provided with water ad libitum and maintained in a 12 h dark/light cycle. The animal work was approved by the UIC Office of Animal Care. Animal Protocol numbers used in this study are ACC 16-180, ACC 19-139, and ACC 20-058-179.

### Induction of breast cancer by DMBA in mice

Female VDR^loxp^ and VDR^ΔIEC^ mice (6-7 week old) were randomly assigned to either control or DMBA groups. Mice were given 6 times weekly with a dose of 1.0 mg DMBA (Sigma-Aldrich, Milwaukee, WI, USA) in 0.2 ml of corn oil or an equal volume of corn oil alone (vehicle) by oral gavage. Then the mice were mated continuously to provide an oscillating hormonal environment and monitored until tumors development. Starting at 12 weeks of age, mice were examined for mammary tumors twice a week. The tumor volume (V) was calculated with caliper measurements using the formulas V= (W^2^ × L)/2 as described before ^65^. The mice were scarified under anesthesia at week 18 or the time when palpable mammary tumors reached a volume of 2 cm^3^. Tumor counts and measurements were performed in a blinded fashion under a stereo-dissecting microscope (Nikon SMZ1000, Melville, NY, USA).

### Butyrate treatment in mice

Female VDR^loxp^ and VDR^ΔIEC^ mice (6-7 week old) were randomly assigned to either DMBA alone or DMBA-butyrate groups. DMBA Control group received filtered drinking water without sodium butyrate. The DMBA-butyrate-treated group received sodium butyrate (Sigma-Aldrich, Milwaukee, WI, USA) at 2.5 % concentration in filtered drinking water. Starting at 12 weeks of age, mice were examined for mammary tumors twice a week. The mice were sacrificed under anesthesia at week 18 or the time when palpable mammary tumors reached a volume of 2 cm^3^.

### Probiotic treatment in mice

Female VDR^loxp^ and VDR^ΔIEC^ mice (6-7 week old) were randomly assigned to either DMBA alone or DMBA-probiotic groups. Mice were daily gavaged with *Lactobacillus plantarum* (1×10^7^ CFU) in 0.1 ml of HBSS or an equal volume of HBSS. Starting at 12 weeks of age, mice were examined for mammary tumors twice a week. The mice were sacrificed under anesthesia at week 18 or the time when palpable mammary tumors reached a volume of 2 cm^3^.

### Intestinal permeability

Fluorescein Dextran (Molecular weight 3 kDa, diluted in HBSS) was gavaged (50 mg/g mouse). Four hours later, mouse blood samples were collected for fluorescence intensity measurement, as previously reported ^66^.

### Hematoxylin and eosin staining

Slides containing mouse colon (proximal or distal colon) sections (5 μm) were deparaffinized in xylene and passed through graded alcohol. They were then stained with hematoxylin and eosin following a previously described method ^63^.

### Western blot analysis and antibodies

Mammary tumors and grossly normal mammary glands from parous age-matched control mice were excised and portions of the tissues were prepared for western blot. Mouse colonic epithelial cells were collected by scraping the tissue from the colon of the mouse, including the proximal and distal regions. The cells were sonicated in lysis buffer (10 mM Tris, pH 7.4, 150 mM NaCl, 1 mM EDTA, 1 mM EGTA, pH 8.0, 1% Triton X-100) with 0.2 mM sodium ortho-vanadate, and protease inhibitor cocktail. The protein concentration was measured using the BioRad Reagent (BioRad, Hercules, CA, USA), and then sonicated. Equal amounts of protein were separated by SDS-polyacrylamide gel electrophoresis, transferred to nitrocellulose, and immunoblotted with primary antibodies. The following antibodies were used: anti-ZO-1 (Invitrogen, 33-9100, Carlsbad, CA, USA), anti-p-β-catenin(552) (Cell Signaling, 9566, Danvers, MA, USA), anti-β-catenin (BD Biosciences, Franklin Lakes, NJ, USA), anti-VDR (Santa Cruz Biotechnology, SC-13133, Dallas, TX, USA), or anti-β-actin (Sigma-Aldrich, A5316, St. Louis, MO, USA) antibodies and were visualized by ECL (Thermo Fisher Scientific, 32106, Waltham, MA, USA). Membranes that were probed with more than one antibody were stripped before re-probing.

### Immunofluorescence

Colonic or breast tissues were freshly isolated and embedded in paraffin wax after fixation with 10% neutral buffered formalin. Immunofluorescence was performed on paraffin-embedded sections (4 μm), after preparation of the slides as described previously ^67^ followed by incubation for 1 hour in blocking solution (2% bovine serum albumin, 1% goat serum in HBSS) to reduce nonspecific background. The tissue samples were incubated overnight with primary antibody ZO-1 at 4°C. Slides were washed 3 times for 5 minutes each at room temperature in wash buffer. Samples were then incubated with the secondary antibody (goat anti-rabbit Alexa Fluor 488, Molecular Probes, CA; 1:200) for 1 hour at room temperature. Tissues were mounted with SlowFade Antifade Kit (Life technologies, s2828, Grand Island, NY, USA), followed by a coverslip, and the edges were sealed to prevent drying. Specimens were examined with a Zeiss laser scanning microscope LSM 710 (Carl Zeiss Inc., Oberkochen, Germany).

### Immunohistochemistry (IHC)

After preparation of the slides, antigen retrieval was achieved by incubating the slides for 15 min in hot preheated sodium citrate (pH 6.0) buffer followed by 30 min of cooling at room temperature. Endogenous peroxidases were quenched by incubating the slides in 3% hydrogen peroxide for 10 min, followed by three rinses with HBSS, and incubation for 1 hour in 3% BSA + 1% goat serum in HBSS to reduce nonspecific background. Primary antibodies VDR or p-β-catenin (552) were applied for overnight in a cold room. After three rinses with HBSS, the slides were incubated in secondary antibody (1:100, Jackson ImmunoResearch Laboratories, Cat.No.115-065-174, West Grove, PA, USA) for 1 hour at room temperature. After washing with HBSS for 10 minutes, the slides were incubated with vectastain ABC reagent (Vector Laboratories, Cat.No. PK-6100, Burlingame, CA 94010, USA) for 1 hour. After washing with HBSS for five minutes, color development was achieved by applying a peroxidase substrate kit (Vector Laboratories, Cat.No. SK-4800, Burlingame, CA 94010) for 2 to 5 minutes, depending on the primary antibody. The duration of peroxidase substrate incubation was determined through pilot experiments and was then held constant for all of the slides. After washing in distilled water, the sections were counterstained with haematoxylin (Leica, Cat.No.3801570, Wetzlar, Germany), dehydrated through ethanol and xylene, and cover-slipped using a permount (Fisher Scientific, Cat.No.SP15-100, Waltham, MA, USA).

### Terminal deoxynucleotidyl transferase dUTP nick end labeling (TUNEL) staining

The number of apoptotic cells were determined using the *In Situ* Cell Death Detection Kit (Sigma-Aldrich, 11684795910, St. Louis, MO, USA) in paraffin-embedded tissue sections. Briefly, antigen retrieval was achieved after deparaffination and rehydration by incubating the slides for 15 min in hot preheated sodium citrate (pH 6.0) buffer. Then the slides were washed with PBS for 10 minutes. After blocking, the slides were incubated with a TUNEL Reaction Mixture for 1 hour at 37°C. The tissues were mounted with SlowFade Antifade Kit (Life technologies, s2828, Grand Island, NY, USA) after 10-minute wash with PBS. The staining was examined with a Zeiss laser scanning microscope LSM 710 (Carl Zeiss Inc., Oberkochen, Germany).

### Real-time PCR measurement of bacterial DNA

DNA was extracted from mouse fecal using EZNA Stool DNA Kit (Omega Bio-tek, Inc. D4015-01, Norcross, GA 30071). The quantitative real-time PCR was conducted using the CFX96 Real-time PCR detection system (Bio-Rad Laboratories, Hercules, CA, USA) and iTaq™ Universal SYBR green supermix (Bio-Rad Laboratories, 1725121, Hercules, CA, USA) according to the manufacturer’s directions. All expression levels were normalized to universal bacteria levels of the same sample. Percent expression was calculated as the ratio of the normalized value of each sample to that of the corresponding untreated control cells. All real-time PCR reactions were performed in triplicate. Primer sequences were designed using Primer-BLAST or were obtained from Primer Bank primer pairs listed in **Table 1**.

**Table 1:**
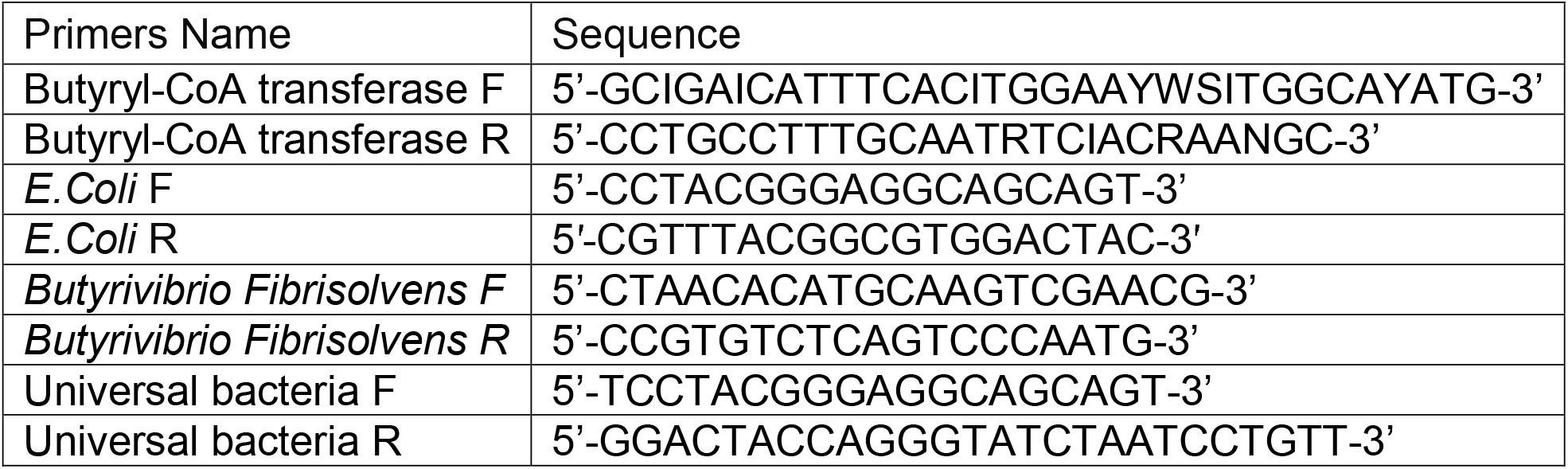
Real-time PCR Primers.

### Multiplex ELISA assay

Mouse blood samples were collected by cardiac puncture and placed in tubes containing EDTA (10 mg/mL). Mouse cytokines were measured using a Cytokine & Chemokine Convenience 26-Plex Mouse ProcartaPlex™ Panel 1 (Invitrogen, EPXR260-26088-90, Carlsbad, CA) according to the manufacturer’s instructions. Briefly, beads of defined spectral properties were conjugated to protein-specific capture antibodies and added along with samples (including standards of known protein concentration, control samples, and test samples) into the wells of a filter-bottom microplate, where proteins bound to the capture antibodies over the course of a 2-hour incubation. After washing the beads, protein-specific biotinylated detector antibodies were added and incubated with the beads for 1 hour. After removal of excess biotinylated detector antibodies, the streptavidin-conjugated fluorescent protein R-phycoerythrin was added and allowed to incubate for 30 minutes. After washing to remove unbound streptavidin–R-phycoerythrin, the beads were analyzed with the Luminex detection system (Bio-rad, Bio-Plex 200 Systems, Hercules, CA).

### Serum LPS detection

LPS in serum samples was measured with limulus amebocyte lysate chromogenic end point assays (Hycult Biotech, HIT302, Plymouth, PA) according to the manufacturer’s indications. The samples were diluted 1:4 with endotoxin-free water and then heated at 75°C for 5 minutes on a warm plate to denature the protein before the reaction. A standard curve was generated and used to calculate the concentrations, which were expressed as EU/mL, in the serum samples.

### Shotgun metagenomic sequencing and Bioinformatic analysis

#### Study design and sampling

VDR^ΔIEC^ and control VDR^LoxP^ mouse strains used in this study were from mice (male and female; aged 6 to 8 weeks). All mice for VDR^LoxP^ (male n=3 and female n=7) and VDR^ΔIEC^ (male n=5 and female n=5) were randomly assigned to each group.

#### Fecal sample collection and shotgun metagenomic sequencing

Fresh fecal sample were collected and placed into the sterile tubes with dry ice and were sent to the University of Illinois at Chicago Research Resources Center for genomic sequencing. The sample DNAs were extracted with DNeasy Power Fecal Kit (Qiagen, Hilden, Germany) according to the manufacturer’s instructions and modified slightly as described previously ^9^. The shotgun metagenomic sequencing was performed with Illumina HiSeq system as described in our previous publications ^12, 68^. After quality checking, filtering the reads, noisy sequences removing, and metagenomic assembling were performed ^69^, the resulting assemblies were filtered. The DNA reads shorter than 1,000 nucleotides were excluded and were classified with Centrifuge ^70^. Finally, each gene was taxonomically annotated by searching for the comprehensive NCBI GenBank non-redundant nucleotide database.

### Statistical Analysis

All data were expressed as the mean ± SD. All statistical tests were 2-sided. All *p*-values < 0.05 were considered statistically significant. Based on data distributions, the differences between samples were analyzed using Welch’s *t*-test or unpaired *t*-test for two groups and one-way ANOVA for more than two groups as appropriate, respectively. Statistical analyses were performed using GraphPad Prism 8 (GraphPad, Inc., San Diego, CA., USA).

## Author Contributions

YZ, JZ: acquisition, analysis, and interpretation of data; drafting the manuscript; and statistical analysis. JZ, SD, SG: assistance with animal models. YX: statistical analysis, and manuscript drafting. JS: study concept and design, analysis and interpretation of data, writing the manuscript for important intellectual content, obtained funding, and study supervision.

## Funding

This research was funded by the UIC Cancer Center, the NIDDK/National Institutes of Health grant R01 DK105118 and R01DK114126, VA Merit Award VA 1 I01 BX004824-01, and DOD BC160450P1 to Jun Sun. The study sponsors played no role in the study design, data collection, analysis, and interpretation of data. The contents do not represent the views of the United States Department of Veterans Affairs or the United States Government.

## Conflicts of Interest

The authors declare no conflict of interest. The funders played no role in the study design, the collection, analyses, or interpretation of data, the writing of the manuscript, or the decision to publish the results.

## Data Availability Statement

The data and materials supporting the results or analyses presented in their paper freely available upon request.

## Notes

### Competing Interest Statement

The authors have declared no competing interest.

## References

1. Shang M, Sun J. Vitamin D/VDR, Probiotics, and Gastrointestinal Diseases. Curr Med Chem 2017; 24:876–87.

2. Sun J. Dietary vitamin D, vitamin D receptor, and microbiome. Current opinion in clinical nutrition and metabolic care 2018; 21:471–4.

3. Abreu MT, Kantorovich V, Vasiliauskas EA, Gruntmanis U, Matuk R, Daigle K, et al. Measurement of vitamin D levels in inflammatory bowel disease patients reveals a subset of Crohn’s disease patients with elevated 1,25-dihydroxyvitamin D and low bone mineral density. Gut 2004; 53:1129–36.

4. Lim WC, Hanauer SB, Li YC. Mechanisms of disease: vitamin D and inflammatory bowel disease. Nat Clin Pract Gastroenterol Hepatol 2005; 2:308–15.

5. Sentongo TA, Semaeo EJ, Stettler N, Piccoli DA, Stallings VA, Zemel BS. Vitamin D status in children, adolescents, and young adults with Crohn disease. The American journal of clinical nutrition 2002; 76:1077–81.

6. Thorne JaC, J. M. The molecular cell biology of VDR. Springer Science & Business Media, 2010.

7. Wu S, Zhang YG, Lu R, Xia Y, Zhou D, Petrof EO, et al. Intestinal epithelial vitamin D receptor deletion leads to defective autophagy in colitis. Gut 2015; 64:1082–94.

8. Wang J, Thingholm LB, Skieceviciene J, Rausch P, Kummen M, Hov JR, et al. Genome-wide association analysis identifies variation in vitamin D receptor and other host factors influencing the gut microbiota. Nat Genet 2016; 48:1396–406.

9. Zhang J, Zhang Y, Xia Y, Sun J. Imbalance of the intestinal virome and altered viral-bacterial interactions caused by a conditional deletion of the vitamin D receptor. Gut Microbes 2021; 13:1957408.

10. Zhang YG, Lu R, Wu S, Chatterjee I, Zhou D, Xia Y, et al. Vitamin D Receptor Protects Against Dysbiosis and Tumorigenesis via the JAK/STAT Pathway in Intestine. Cell Mol Gastroenterol Hepatol 2020; 10:729–46.

11. Zhang Y, Garrett S, Carroll RE, Xia Y, Sun J. Vitamin D receptor upregulates tight junction protein claudin-5 against colitis-associated tumorigenesis. Mucosal Immunol 2022.

12. Lu R, Zhang YG, Xia Y, Zhang J, Kaser A, Blumberg R, et al. Paneth Cell Alertness to Pathogens Maintained by Vitamin D Receptors. Gastroenterology 2021; 160:1269–83.

13. Wu S, Liao AP, Xia Y, Li YC, Li JD, Sartor RB, et al. Vitamin D receptor negatively regulates bacterial-stimulated NF-kappaB activity in intestine. Am J Pathol 2010; 177:686–97.

14. Fukata M, Abreu MT. Pathogen recognition receptors, cancer and inflammation in the gut. Curr Opin Pharmacol 2009; 9:680–7.

15. Bourzac KM, Guillemin K. Helicobacter pylori-host cell interactions mediated by type IV secretion. Cellular microbiology 2005; 7:911–9.

16. Bourzac KM, Satkamp LA, Guillemin K. The Helicobacter pylori cag pathogenicity island protein CagN is a bacterial membrane-associated protein that is processed at its C terminus. Infection and immunity 2006; 74:2537–43.

17. Gobert AP, McGee DJ, Akhtar M, Mendz GL, Newton JC, Cheng Y, et al. Helicobacter pylori arginase inhibits nitric oxide production by eukaryotic cells: a strategy for bacterial survival. Proc Natl Acad Sci U S A 2001; 98:13844–9.

18. Yan F, Cao H, Chaturvedi R, Krishna U, Hobbs SS, Dempsey PJ, et al. Epidermal growth factor receptor activation protects gastric epithelial cells from Helicobacter pylori-induced apoptosis. Gastroenterology 2009; 136:1297–307, e1-3.

19. Terzic J, Grivennikov S, Karin E, Karin M. Inflammation and colon cancer. Gastroenterology 2010; 138:2101–14 e5.

20. Akram M, Iqbal M, Daniyal M, Khan AU. Awareness and current knowledge of breast cancer. Biological research 2017; 50:33.

21. Iqbal MUN, Khan TA. Association between Vitamin D receptor (Cdx2, Fok1, Bsm1, Apa1, Bgl1, Taq1, and Poly (A)) gene polymorphism and breast cancer: A systematic review and meta-analysis. Tumour biology : the journal of the International Society for Oncodevelopmental Biology and Medicine 2017; 39:1010428317731280.

22. Atoum M, Alzoughool F. Vitamin D and Breast Cancer: Latest Evidence and Future Steps. Breast cancer : basic and clinical research 2017; 11:1178223417749816.

23. Zhang YG, Wu S, Sun J. Vitamin D, Vitamin D Receptor, and Tissue Barriers. Tissue Barriers 2013; 1.

24. Song M, Chan AT, Sun J. Influence of the Gut Microbiome, Diet, and Environment on Risk of Colorectal Cancer. Gastroenterology 2020; 158:322–40.

25. Richardson AL, Wang ZC, De Nicolo A, Lu X, Brown M, Miron A, et al. X chromosomal abnormalities in basal-like human breast cancer. Cancer Cell 2006; 9:121–32.

26. González-Beiras C, Marks M, Chen CY, Roberts S, Mitjà O. Epidemiology of Haemophilus ducreyi Infections. Emerging infectious diseases 2016; 22:1–8.

27. Luo S, Yin J, Peng Y, Xie J, Wu H, He D, et al. Glutathione is Involved in Detoxification of Peroxide and Root Nodule Symbiosis of Mesorhizobium huakuii. Current microbiology 2020; 77:1–10.

28. Komaniecka I, Zdzisinska B, Kandefer-Szerszen M, Choma A. Low endotoxic activity of lipopolysaccharides isolated from Bradyrhizobium, Mesorhizobium, and Azospirillum strains. Microbiology and immunology 2010; 54:717–25.

29. Holeček M. Histidine in Health and Disease: Metabolism, Physiological Importance, and Use as a Supplement. Nutrients 2020; 12.

30. Kulis-Horn RK, Persicke M, Kalinowski J. Histidine biosynthesis, its regulation and biotechnological application in Corynebacterium glutamicum. Microbial biotechnology 2014; 7:5–25.

31. Hudspeth DS, Vary PS. spoVG sequence of Bacillus megaterium and Bacillus subtilis. Biochim Biophys Acta 1992; 1130:229–31.

32. Zhang J, Xia Y, Sun J. Breast and gut microbiome in health and cancer. Genes Dis 2021; 8:581–9.

33. Yang J, Tan Q, Fu Q, Zhou Y, Hu Y, Tang S, et al. Gastrointestinal microbiome and breast cancer: correlations, mechanisms and potential clinical implications. Breast Cancer 2017; 24:220–8.

34. Soto-Pantoja DR, Gaber M, Arnone AA, Bronson SM, Cruz-Diaz N, Wilson AS, et al. Diet Alters Entero-Mammary Signaling to Regulate the Breast Microbiome and Tumorigenesis. Cancer Res 2021; 81:3890–904.

35. Buchta Rosean C, Bostic RR, Ferey JCM, Feng TY, Azar FN, Tung KS, et al. Preexisting Commensal Dysbiosis Is a Host-Intrinsic Regulator of Tissue Inflammation and Tumor Cell Dissemination in Hormone Receptor-Positive Breast Cancer. Cancer Res 2019; 79:3662–75.

36. Zhu J, Liao M, Yao ZT, Liang WY, Li QB, Liu JL, et al. Breast cancer in postmenopausal women is associated with an altered gut metagenome. Microbiome 2018; 6.

37. Huggins C, Grand LC, Brillantes FP. Mammary cancer induced by a single feeding of polymucular hydrocarbons, and its suppression. Nature 1961; 189:204–7.

38. Ethier SP, Ullrich RL. Induction of mammary tumors in virgin female BALB/c mice by single low doses of 7,12-dimethylbenz[a]anthracene. J Natl Cancer Inst 1982; 69:1199–203.

39. Currier N, Solomon SE, Demicco EG, Chang DL, Farago M, Ying H, et al. Oncogenic signaling pathways activated in DMBA-induced mouse mammary tumors. Toxicol Pathol 2005; 33:726–37.

40. Abba MC, Zhong Y, Lee J, Kil H, Lu Y, Takata Y, et al. DMBA induced mouse mammary tumors display high incidence of activating Pik3caH1047 and loss of function Pten mutations. Oncotarget 2016; 7:64289–99.

41. Lee G, Goretsky T, Managlia E, Dirisina R, Singh AP, Brown JB, et al. Phosphoinositide 3-kinase signaling mediates beta-catenin activation in intestinal epithelial stem and progenitor cells in colitis. Gastroenterology 2010; 139:869–81, 81 e1-9.

42. Wang S, Han Y, Zhang J, Yang S, Fan Z, Song F, et al. Me6TREN targets beta-catenin signaling to stimulate intestinal stem cell regeneration after radiation. Theranostics 2020; 10:10171–85.

43. Perry JM, Tao F, Roy A, Lin T, He XC, Chen S, et al. Overcoming Wnt-beta-catenin dependent anticancer therapy resistance in leukaemia stem cells. Nat Cell Biol 2020; 22:689–700.

44. Zeng MY, Inohara N, Nunez G. Mechanisms of inflammation-driven bacterial dysbiosis in the gut. Mucosal Immunol 2017; 10:18–26.

45. Louis P, Duncan SH, McCrae SI, Millar J, Jackson MS, Flint HJ. Restricted distribution of the butyrate kinase pathway among butyrate-producing bacteria from the human colon. J Bacteriol 2004; 186:2099–106.

46. Wu SP, Yoon S, Zhang YG, Lu R, Xia YL, Wan JD, et al. Vitamin D receptor pathway is required for probiotic protection in colitis. Am J Physiol-Gastr L 2015; 309:G341–G9.

47. Zhang J, Lu R, Zhang Y, Matuszek Z, Zhang W, Xia Y, et al. tRNA Queuosine Modification Enzyme Modulates the Growth and Microbiome Recruitment to Breast Tumors. Cancers (Basel) 2020; 12.

48. De Mattia E, Cecchin E, Montico M, Labriet A, Guillemette C, Dreussi E, et al. Association of STAT-3 rs1053004 and VDR rs11574077 With FOLFIRI-Related Gastrointestinal Toxicity in Metastatic Colorectal Cancer Patients. Front Pharmacol 2018; 9:367.

49. Fu A, Yao B, Dong T, Chen Y, Yao J, Liu Y, et al. Tumor-resident intracellular microbiota promotes metastatic colonization in breast cancer. Cell 2022; 185:1356–72 e26.

50. Parida S, Wu S, Siddharth S, Wang G, Muniraj N, Nagalingam A, et al. A Procarcinogenic Colon Microbe Promotes Breast Tumorigenesis and Metastatic Progression and Concomitantly Activates Notch and beta-Catenin Axes. Cancer Discov 2021; 11:1138–57.

51. Urbaniak C, Gloor GB, Brackstone M, Scott L, Tangney M, Reid G. The Microbiota of Breast Tissue and Its Association with Breast Cancer. Applied and environmental microbiology 2016; 82:5039–48.

52. Sun J, Zhang YG. Vitamin D Receptor Influences Intestinal Barriers in Health and Disease. Cells 2022; 11.

53. Soldatenkov VA, Prasad S, Voloshin Y, Dritschilo A. Sodium butyrate induces apoptosis and accumulation of ubiquitinated proteins in human breast carcinoma cells. Cell death and differentiation 1998; 5:307–12.

54. Protiva P, Cross HS, Hopkins ME, Kallay E, Bises G, Dreyhaupt E, et al. Chemoprevention of colorectal neoplasia by estrogen: potential role of vitamin D activity. Cancer Prev Res (Phila Pa) 2009; 2:43–51.

55. Murillo G, Mehta RG. Chemoprevention of chemically-induced mammary and colon carcinogenesis by 1alpha-hydroxyvitamin D5. J Steroid Biochem Mol Biol 2005; 97:129–36.

56. Kaler P, Augenlicht L, Klampfer L. Macrophage-derived IL-1beta stimulates Wnt signaling and growth of colon cancer cells: a crosstalk interrupted by vitamin D3. Oncogene 2009; 28:3892–902.

57. Fichera A, Little N, Dougherty U, Mustafi R, Cerda S, Li YC, et al. A vitamin D analogue inhibits colonic carcinogenesis in the AOM/DSS model. J Surg Res 2007; 142:239–45.

58. Nagpal S, Lu J, Boehm MF. Vitamin D analogs: mechanism of action and therapeutic applications. Curr Med Chem 2001; 8:1661–79.

59. Palmer HG, Sanchez-Carbayo M, Ordonez-Moran P, Larriba MJ, Cordon-Cardo C, Munoz A. Genetic signatures of differentiation induced by 1alpha,25-dihydroxyvitamin D3 in human colon cancer cells. Cancer Res 2003; 63:7799–806.

60. Chan AT, Giovannucci EL. Primary prevention of colorectal cancer. Gastroenterology 2010; 138:2029–43 e10.

61. Gombart AF, Luong QT, Koeffler HP. Vitamin D compounds: activity against microbes and cancer. Anticancer Res 2006; 26:2531–42.

62. Barrett JC, Hansoul S, Nicolae DL, Cho JH, Duerr RH, Rioux JD, et al. Genome-wide association defines more than 30 distinct susceptibility loci for Crohn’s disease. Nat Genet 2008; 40:955–62.

63. Zhang Y-g, Lu R, Xia Y, Zhou D, Petrof E, Claud EC, et al. Lack of Vitamin D Receptor Leads to Hyperfunction of Claudin-2 in Intestinal Inflammatory Responses. Inflammatory Bowel Diseases 2018; 25:97–110.

64. Zhang Y-G, Lu R, Wu S, Chatterjee I, Zhou D, Xia Y, et al. Vitamin D receptor protects against dysbiosis and tumorigenesis via the JAK/STAT pathway in intestine. Biorxiv 2020; 02.

65. Faustino-Rocha A, Oliveira PA, Pinho-Oliveira J, Teixeira-Guedes C, Soares-Maia R, da Costa RG, et al. Estimation of rat mammary tumor volume using caliper and ultrasonography measurements. Lab animal 2013; 42:217–24.

66. Zhang YG, Lu R, Xia Y, Zhou D, Petrof E, Claud EC, et al. Lack of Vitamin D Receptor Leads to Hyperfunction of Claudin-2 in Intestinal Inflammatory Responses. Inflamm Bowel Dis 2019; 25:97–110.

67. Lu R, Wu S, Liu X, Xia Y, Zhang YG, Sun J. Chronic effects of a Salmonella type III secretion effector protein AvrA in vivo. PLoS One 2010; 5:e10505.

68. Chatterjee I, Lu R, Zhang Y, Zhang J, Dai Y, Xia Y, et al. Vitamin D receptor promotes healthy microbial metabolites and microbiome. Scientific Reports 2020; 10:7340.

69. Xia Y, Sun J, Chen D-G. Bioinformatic analysis of microbiome data. Statistical Analysis of Microbiome Data with R: Springer, 2018:1–27.

70. Kim D, Song L, Breitwieser FP, Salzberg SL. Centrifuge: rapid and sensitive classification of metagenomic sequences. Genome research 2016; 26:1721–9.

